# Constraint-based modeling identifies metabolic vulnerabilities during the epithelial to mesenchymal transition

**DOI:** 10.1101/2022.01.31.478483

**Authors:** Scott Campit, Venkateshwar G. Keshamouni, Sriram Chandrasekaran

## Abstract

Epithelial-to-mesenchymal transition (EMT) is a developmentally conserved cellular process critical for tumor metastasis. EMT enables malignant epithelial cells to acquire mesenchymal-like migratory and invasive phenotype. During EMT cancer cells undergo extensive metabolic reprogramming that correlates with the suppression of proliferation, and stimulation of the energy-intensive migratory behavior. However, the causal relationship between metabolic changes and coordinated physiological phenotypes that occur during EMT is still unclear. We used bulk time-course transcriptomics and proteomics, and single-cell transcriptomics from five independent EMT studies in A549 lung adenocarcinoma cells to simulate metabolic network activity using constraint-based modeling. Model predictions were validated using literature mining, experimental studies and CRISPR-Cas9 essentiality screens.

We uncovered temporal metabolic dependencies in glycolysis and glutamine metabolism reactions, and experimentally validated isoform-specific dependency on Enolase3 for cell survival during EMT. Together, our approach uncovered temporally regulated cell-state-specific metabolic dependencies in cells undergoing EMT.

## Introduction

EMT is a reversible developmental process stimulated by extracellular signals that facilitate the transition from an epithelial (E) cell to a motile and invasive mesenchymal-like (M) cells, enabling circulating tumor cells to initiate metastasis (Pastushenko and Blanpain 2019). More importantly, EMT is not a binary process but occurs through a spectrum of distinct intermediate states with potential functional consequences. The cytokine - transforming growth factor **β** (TGF-**β**), is a potent inducer of EMT. Consistently, TGF- **β** levels are highly upregulated and directly correlate with tumor progression, enhanced invasion, metastasis, and poor survival in patients with non-small cell lung cancer (NSCLC) (Padua and Massagué 2009). Cell culture models of TGF-**β** -induced EMT serves as robust *in vitro* model to investigate mechanisms of metastasis. In addition to metastasis, the process of EMT-MET is implicated in several clinically relevant aspects including, tumor heterogeneity, stemness, and drug resistance (Ramesh et al. 2020). Understanding underlying regulatory mechanisms is essential to develop therapeutic strategies that can prevent EMT or promote MET to inhibit metastasis.

Extensive molecular and structural changes that occur during EMT can potentially induce robust metabolic reprogramming to support changing cellular phenotypes. Cancers show enhanced glycolytic pathway activity even in the presence of oxygen, via the Warburg effect (Vander Heiden et al. 2009). It is now clear that the Warburg effect is not restricted to cancer cells but it is an adaptive physiological program that occurs in many normal cell types, when cells need rapid ATP production, biomass synthesis and balancing of reactive oxygen species (Pålsson-McDermott and O’Neill 2020)(Kim et al. 2016), Studies have demonstrated an enhanced glycolytic activity during EMT. However, similar to EMT, metabolic reprogramming may also involve a spectrum of pathway combinations at a given cellular steady state. While several metabolic rewiring strategies have been observed in diverse cancers, a comprehensive systems-level characterization of metabolic reprogramming during the EMT has not been carried out. Methods that can infer metabolic dysregulation using omics data will be invaluable for understanding causal relationship between EMT and metabolic reprogramming.

Constraint-Based Optimization and Reconstruction Analysis (COBRA) is a widely used approach for simulating genome-scale metabolic fluxes using omics data. COBRA simulates metabolic fluxes by using the metabolic network architecture, nutrient availability, and omics data as constraints in an optimization problem representing a cellular objective, such as maximizing biomass production (Orth et al. 2010). COBRA models have inferred metabolic rewiring strategies in several cancer subtypes (Oruganty et al. 2020; Yizhak et al. 2014; Nilsson et al. 2020). For instance, incorporating metabolomics data to identify synthetically lethal metabolic genes in pancreatic cancer (Nelson et al. 2020). However, to our knowledge, no one has applied COBRA to study metabolic network heterogeneity during the EMT and characterized the metabolic properties of intermediate states during EMT.

We used COBRA to simulate metabolic activity and vulnerabilities during EMT using diverse omics sources, including time-course transcriptomics, proteomics, and single-cell transcriptomics data. Notably, this study applies constraint-based modeling to single-cell cancer transcriptomics data to capture the metabolic heterogeneity during EMT. From our analysis, we were able to identify known metabolic dependencies during EMT, such as uptake of glucose and glutamine. We also predicted new metabolic dependencies including the enolase and GOT1 reactions and those related to alpha-ketoglutarate metabolism. Surprisingly, many of these dependencies were time-specific, suggesting that there is a narrow temporal window during which the cells can be targeted with drugs that inhibit these pathways. We also found metabolic changes that showed consistent trends based on model predictions derived from both the bulk and single-cell studies. Together, our analysis provides a framework to integrate multiple omics datasets to examine tumor metabolic heterogeneity and infer new drug targets.

## Methods

### Differential Expression (Bulk Studies)

We analyzed two transcriptomics (Hecker et al. 2009; Keshamouni et al. 2009) and two proteomics (Keshamouni et al. 2006; Lu et al. 2019) EMT time-course studies with A549 as the cell model undergoing TGF-*β* induction. All studies compared later time point after TGF-*β* induction over day 0 to obtain differentially expressed genes and proteins. When possible, authors’ methods and provided datasets were used to obtain a list of up- and downregulated gene sets. If no preprocessed data was provided (as in the case of GSE17518), *limma-voom* (Law et al. 2014) was performed to determine differentially expressed genes between conditions. Additionally, a GAM-LOESS model was used to determine differentially expressed genes in GSE147405 (Cook and Vanderhyden 2020), aggregating single-cells at the time-course level. The regression coefficients from the GAM-LOESS model were used to determine the sign of regulation (up/down). P-values from *limma-voom* and a GAM-LOESS model were adjusted using the Benjamini-Hochberg method and the significance threshold used was P-value < 0.05. The expression matrix containing statistically significant normalized scores for all metabolic genes across all 5 experiments can be found in **Supplementary Table 1**.

### Individual Cell Differential Expression

We computed differentially expressed genes for individual cells without TGF-*β* removal in GSE147405 to simulate individual cell fluxes and reaction knockout growth rates. Data preprocessing included data scaling, removing contaminant artifacts such as mitochondrial genes, and removing cells with low total gene counts. This was performed on the raw data object. Further, we used the data imputation algorithm MAGIC (Van Dijk, D. et al., 2018) to fill in drop out values. The MAGIC-imputed data was transformed to a Z-score using a Z-score method that subtracts out the median and centers the data based on the median absolute deviation (MAD). The formula for the robust Z-score for a specific gene *i* in a given cell *j* is shown in **Equation 1**:

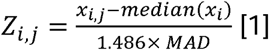

Where 1.486 is a scaling constant. For any given cell, a gene was determined to be upregulated/downregulated if the robust Z-score was positive/negative and the P-value < 0.05.

### Prioritizing metabolic gene targets across multiple studies

We evaluated the robustness metabolic gene dysregulation across five EMT studies using the following prioritization score *η*. The method to compute *η* shown in **Equation 2**:

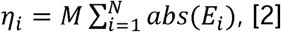

Where *M*is the number of studies where the gene was determined to be significant and *E*_*i*_is the gene effect size (log_2_ fold change or Z-score) for gene *i*. The prioritization scores were ranked in descending order and used to prioritize reactions for further investigation.

### COnstraint-Based Reconstruction and Analysis (COBRA)

Flux balance analysis (FBA; Orth et al. 2010) was used to simulate metabolic activity using the human metabolic reconstruction RECON1 (Duarte et al. 2007). Cells were assumed to maximize biomass production as the objective function. Differentially expressed metabolic genes that intersected with RECON1 were used as biological constraints to maximize (upregulation) or minimize (downregulation) metabolic flux using a modified form of the iMAT algorithm (Zur et al. 2010; Shen et al 2019). Parsimonious enzyme usage (Lewis et al. 2010) was an additional assumption to obtain a unique metabolic flux distribution and to minimize fluxes that did not contribute to biomass formation. Metabolic fluxes and growth rates from single gene and reaction knockout simulations were obtained using COBRA. To ensure a feasible growth rate was calculated, we removed genes/reactions that were upregulated in the knockout and set the percent knockout to be 99% to promote a feasible flux solution.

### Differential Metabolic Activity and Knockout Sensitivity Analysis

To determine differentially active metabolic reactions, we used the priority score described in equation 1 on the absolute value of the metabolic fluxes. Most reactions show zero flux, and so reactions that showed metabolic activity were considered to be overactive metabolic reactions.

To determine the impact metabolic genes have on the growth rate during different stages of EMT, we computed a sensitivity score *θ* comparing EMT versus control growth rates for each gene knockout. The equation to compute the bulk sensitivity score is shown in **Equation 3**:

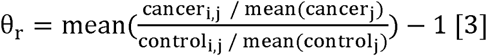

The flux or growth rate was mean-normalized for the control (day 0) and the TGF-*β* treatment (all other days). Then, the final score was taken as the ratio of the TGF-*β* treated growth rate over the average control growth rate for a given reaction knockout. The average ratio across all cells was taken as the score to identify differentially sensitive metabolic reactions. The score was centered at 0. The intuition behind *θ* is as follows: if the score is 0, the gene/reaction knockout has no difference between cancer and control. If the score is less than 0, the knockout impacts the cancer cell more than control, and is considered to be essential for cellular growth. To rank and prioritize metabolic targets for experimental validation, we used the same prioritization score as we did to rank differentially expressed genes.

### Classifying cancer cell line states

Cancer cell lines from the Cancer Cell Line Encyclopedia (CCLE) were annotated by their source from a primary tumor or metastatic tumor. To also classify whether a cell was in the epithelial versus mesenchymal state, we calculated the Z-score and p-value for all genes in the CCLE and mapped them to known EMT markers. Upregulated genes were classified as having a positive Z-score and a significance threshold of p-value < 0.05, while downregulated genes were classified as having a negative Z-score with the same significance threshold.

EMT markers (with up- and downregulated signatures) were taken from MSigDB (Liberzon et al. 2015) across three studies from different tissues of origin induced using TGF-*β*. We further filtered this list with NSCLC markers without up/downregulation annotations from EMTome (Vasaikar et al. 2021). The final number of markers obtained for EMT was 14 genes, which were used to classify cancer cell lines (CCLs). We chose to classify cell lines using upper and lower quantiles of gene makers. Up- and downregulated genes from NSCLC CCLs were cross referenced to the list of EMT signatures and labeled as E if the number of signatures was less than 5 genes or M if the number of signatures was greater than 8.

### CRISPR-Cas9 Analysis

We analyzed batch corrected CERES Scores (Pacini et al. 2021) for metabolic genes that were predicted by COBRA to have increased metabolic activity or resulted in a reduction of growth rate from knockout. CERES Scores were separated based on their association with metastatic (Met) and primary (Prim) cell lines, which was determined based on the CCLE metadata (Barretina et al. 2012). We also compared the CCLE annotations against our own Epithelial (Epi) and Mesenchymal (Mes) annotations, methods described above.

To evaluate how well our model predictions related to CERES Scores, we calculated the Pearson correlation coefficient between our predicted growth scores and ratios of Mes / Epi or Met / Prim CERES Scores. Further, we compared the growth scores against different subsets of the CERES Score data, including NSCLC only cell lines and all cell lines.

### Identifying metabolic enzymes and EMT studies for systematic literature validation

To be considered for our systematic literature validation, we pooled a list of metabolic enzymes predicted from COBRA from bulk and single-cell reaction knockouts that had lethal reactions in at least 2 studies (growth score < 0). The query was performed using PubMed and Google with the following keywords using AND filtering: “EMT”, “Metabolism”, “A549”, “metastasis”, “cancer”, “cancer metabolism”, and the individual gene of interest. The list of the manually curated results can be found in **Supplementary Table 5**.

We expanded the scope of our literature search to encompass all cancer cell lines. The query was manually curated to either support or refute the COBRA predictions. None of the model predictions contradicted the literature. Reaction predictions and their confidence were scored (1-3), where 1 has no evidence based on literature and 3 has strong A549 or lung adenocarcinoma specific evidence. The rules to assign each score for each reaction prediction are shown below:

- **1:** Prediction has no literature support.
- **2:** Prediction has literature evidence with general cancer lineages.
- **3:** Prediction has literature evidence either with specific experiments from A549 or related lung adenocarcinoma tissue/cell lines.

### Cell culture, siRNA transfection and EMT induction

A549 human lung adenocarcinoma cell line was obtained from the American Type Culture Collection (Manassas, VA) and maintained in RPMI-1640 medium with glutamine supplemented with 10% FBS, penicillin, and streptomycin at 37°C in 5% CO2. For inducing EMT cells at 40-50% confluency in complete medium were serum starved for 24 hrs and treated with TGF- *β* (5 ng/ml) for 72 hrs.

Isoform specific siRNA for enolases includes a pool of 4 SMART selection-designed synthetic duplexes (Dharmocon’ s SMARTpool). A scrambled sequence from the same company is used as a control. Cells at 40-50% confluency were transfected with siRNA using Lipofectamine 2000 (Cat No: 18324-012, Invitrogen) and optiMEM medium (Cat No: 31985, Gibco) following the manufacturer’s instructions. After 6 hours of transfection cells were washed and allowed to recover from transfection in RPMI 1640 medium with 10% FBS before inducing EMT as described above.

### Apoptosis assays

Apoptosis was assessed by two independent methods; *1) AnnexinV/7-AAD staining (Kit from Biolegend Cat# 640922):* At the end of the EMT experiment described above, all cells (including floating cells) were collected, washed and resuspended in Annexin V binding buffer. 100 ul of cell suspension was stained with 5 ul of FITC-Annexin V, followed by 5 ul of 7-AAD staining solution. After 30 min incubation at room temperature in dark, 400 ul of Annexin V binding buffer is added and assessed for Annexin V and 7-AAD staining by flow cytometry. Both Annexin V postive (early apoptotic) and Annexin V and 7-AAD double positive cells (late apoptotic) are added together for assessing total apoptosis. 2*) Assessing Caspase 3 activation:* To assess casapase3 activation during EMT, an artificial caspase3 substrate coupled to a green fluorescent DNA-binding dye (DEVD-Nucview) is added to the cell culture. When caspase3 is activated, it cleaves the DNA-binding dye which enters the nuclei and labels an apoptotic cell with green fluorescence allowing its imaging. Green fluorescent apoptotic cells were imaged under a fluorescent microscope 48 hrs after TGF-*β* -induced EMT.

## Results

### COBRA reveals that cells undergoing EMT exhibit enhanced glycolysis during early and late stages

We performed a meta-analysis of differentially expressed genes and proteins across four bulk EMT datasets. Two were RNASeq-based datasets (GSE17708 and GSE17518), and two were proteomics-based datasets (Garcia and Keshamouni). To aggregate the results from multiple studies, we designed a prioritization score to rank the reactions based on effect size and whether or not the gene was significantly expressed in a given study (**Supplementary Table 1; Methods**).

We simulated the metabolic fluxes for each time-point using the transcriptomics and proteomics data to see how metabolic activity changes over time during EMT using Flux Balance Analysis (FBA; **see Methods**). FBA uses a linear optimization procedure with biological constraints, such as knowledge of the metabolic network structure (known as a stoichiometric matrix) and expression levels as inputs to generate cell-state specific metabolic flux profiles (O’Brien et al. 2015). FBA assumes that the cell is maximizing an objective, usually its biomass production. While standard FBA outputs multiple flux profiles due to the rank deficiency of the S matrix (Orth et al. 2010), Parsimonious FBA or pFBA provides a unique flux distribution by assuming optimal enzyme efficiency by minimizing the overall metabolic flux throughout the metabolic network while maximizing biomass production (Lewis et al. 2010). pFBA identifies the smallest set of active reactions that best support biomass production.

Our predictions using pFBA reveal that there are more active reactions during the early and late phases of EMT. During the intermediate phases of EMT, metabolic activity goes down. As cells undergo dramatic structural rearrangements when transitioning to a mesenchymal cell, cells require energetic substrates such as ATP to facilitate these processes. Our metabolic model assumed that these cancer cells were optimizing for increased biomass production, and the reduction of fluxes for biomass production during the intermediate EMT stages suggests that metabolic activity is being siphoned towards other processes such as motility. Our metabolic flux profile data suggests that cancer cells upregulate metabolism initially to build up metabolic substrate levels, and then divert all transcriptional resources towards other processes.

Samples within these time-points tend to have similar metabolic functions, as most active reactions are found within central carbon metabolism (glycolysis/gluconeogenesis, pentose phosphate pathway, folate metabolism) and nutrient exchange subsystems. These metabolic pathways contribute to biomass formation. We visualized the top 50 reactions sorted by prioritization scores (**Methods; S. Figure 1; Supplementary Table 2**). The prioritization score takes into consideration the number of studies where a given metabolic gene(s) encoding a reaction was determined to be significant and absolute value of the gene effect size (log_2_ fold change or Z-score). Developing a prioritization score enabled us to filter through 3744 reactions to provide a concise reaction list for downstream analyses.

Several glycolytic reactions were predicted to have increased metabolic activity and priority scores (**Figure 1A**), which was expected given how cancer cells rewire glycolytic activity, as evidenced by the Warburg effect (Vander Heiden et al. 2009). Several glycolytic substrates play a role in both cellular survival and cancer proliferation. It has been well established that TGF-*β* increases expression of several glycolytic enzymes (Jia et al. 2021). We found that hexokinase, glyceraldehyde-3-phosphate dehydrogenase (GAPDH), and enolase were the top 3 glycolytic reactions that were highly active in both early and late EMT stages, supporting previous studies that suggest glycolysis is directly impacted by TGF-*β* induction. The timing of metabolic activity suggests that glycolysis is essential for initiating EMT and establishing metastasis at later stages.

**Figure 1.**
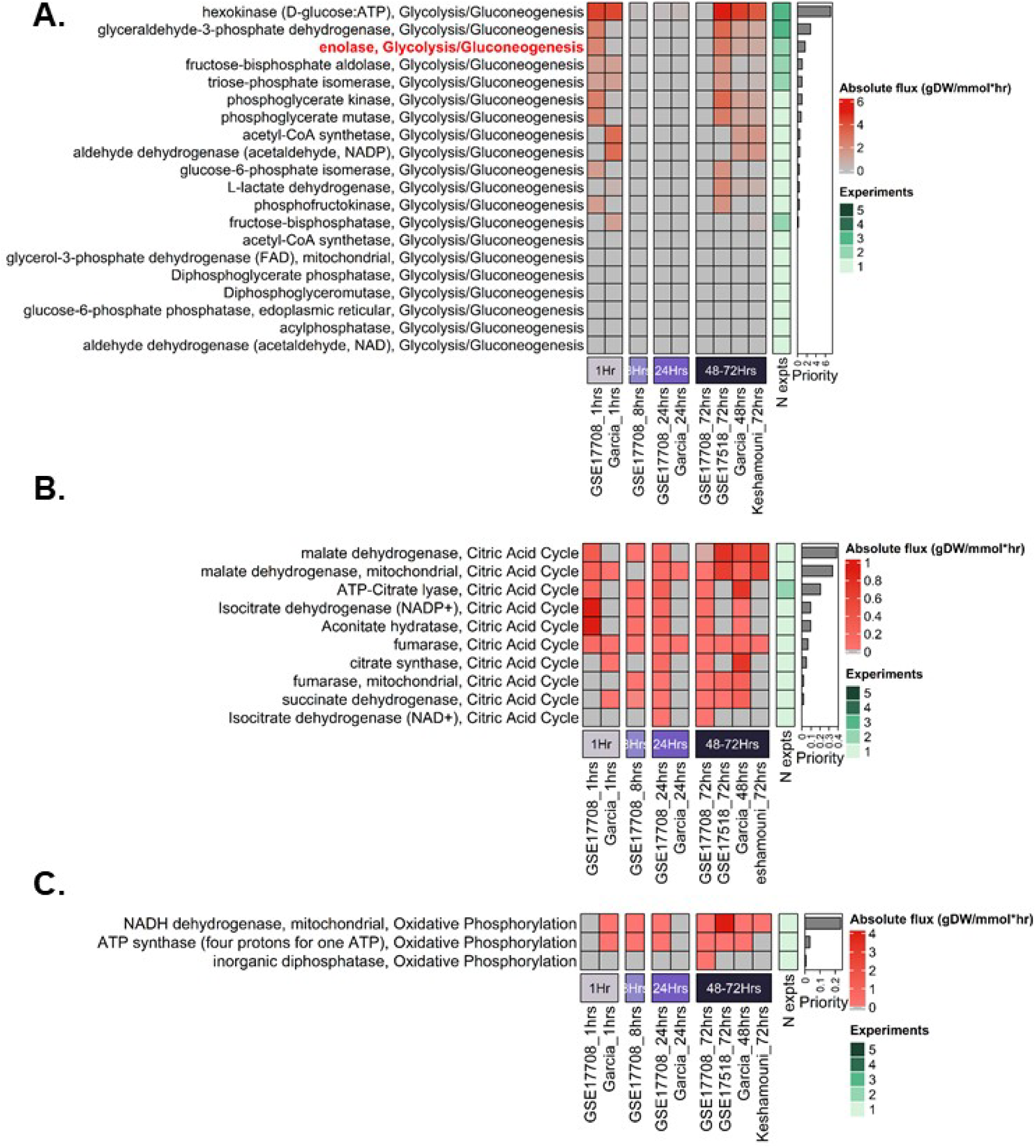
A549 metabolism is predicted to be overactive during the early and late phases of EMT. A. Reactions are sorted based on the priority score, which is a function of the number of studies with significant metabolic genes that encode the reaction and the effect size (Z-score or log2 fold change). The metabolic fluxes were simulated using the RECON1 human metabolic reconstruction. Several metabolic reactions within the Glycolysis/Gluconeogenesis metabolic subsystem are overactive in the earlier stages (1hr) and late stages (48-72hrs) of EMT, based on the absolute value of the metabolic fluxes predicted by constraint-based modeling. The top 5 reactions in the Glycolysis/Gluconeogenesis subsystem have at least two studies supporting the flux predictions. Enolase is bolded as it was prioritized for experimental validation. B. Metabolic reactions within the Citric Acid Cycle are predicted to have more uniform activity across all time points relative to control (unconstrained flux distribution) and have lower priority scores compared to glycolysis. C. Metabolic reactions within the Oxidative Phosphorylation metabolic subsystem are also predicted to have more uniform activity relative to control across all time points.

### Genome-scale reaction knockout simulation identifies extensive vulnerabilities in mesenchymal state

While our previous analysis focused on reaction fluxes, next we used FBA to simulate the impact of reaction knockout on cellular growth in each time-points across five independent A549 TGF-*β* induced EMT studies (**S. Figure 2**; **Supplementary Table 3**). Briefly, each metabolic reaction encoded in the reconstruction was systematically shut off (upper and lower bounds were set to 0) to simulate a “knockout”, while the growth rate objective was optimized. This method allows us to infer the impact systematic reaction knockouts have on cellular growth. We analyzed the distribution of knockouts across bulk experiments and by time-course. The later stage of EMT were predicted to have more vulnerabilities (932 reactions) than in the early stage (874 reactions) and intermediate stage (660 reactions), suggesting mesenchymal cells are more vulnerable to metabolic perturbation (**Figure 2; inset**).

**Figure 2.**
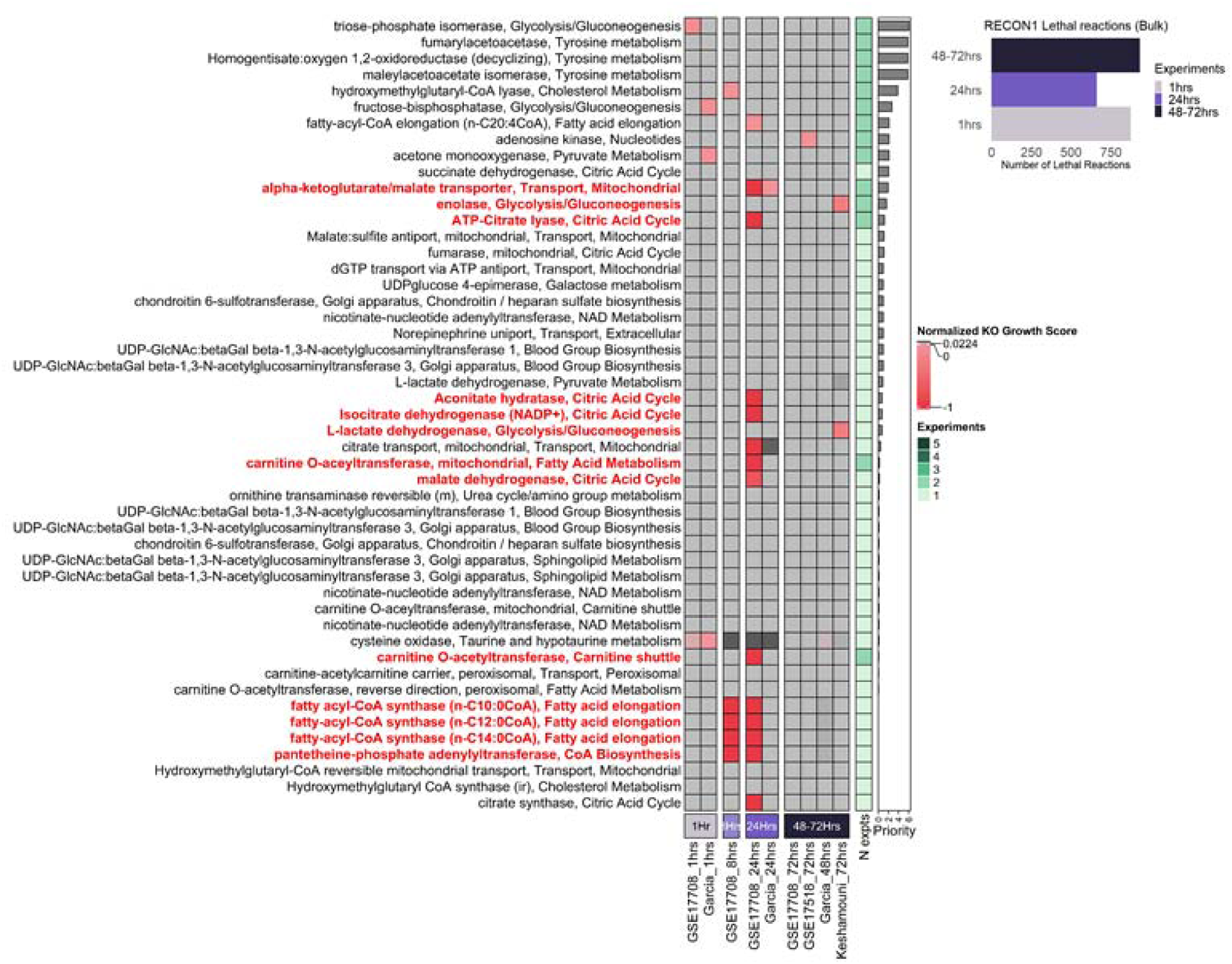
A global view of cancer growth sensitivity to metabolic reaction knockout during EMT. Normalized KO Growth Scores closer to a value of -1 confer a decrease in growth relative upon simulated reaction knockout relative to the control (an unconstrained metabolic reconstruction growth rate). Reactions are sorted based on priority scores. The barplots (inset) show the distribution of lethal reactions (Growth Score < 0) for three timepoints 1 hour, 24 hours, and 48-72 hours after TGF**-β**induction across all experiments in 3 time points. Reactions with red and bold text were predicted to be highly sensitive to knockout and were prioritized for downstream analyses with CERES essentiality scores.

We identified over 40 reactions that were specifically sensitive in specific EMT stages and studies (**Figure 2**). These also highlight the technical and biological variance that is observed in EMT studies across different omics modes. Notably, alpha-ketoglutarate (AKG) transport between the cytosol was unique to the intermediate stage in two out of five independent studies. From a mechanistic standpoint, AKG likely suppresses metastasis by counteracting the effects of other oncometabolites such as 2-hydroxyglutarate, succinate, and fumarate (Wei et al. 2020). While the impact of AKG and cellular differentiation / proliferation has been observed through several nutrient perturbation studies in cancer and stem-cells, the exact source and subcellular contribution of AKG and its impact on metastasis is difficult to determine experimentally. Our computational model suggests that knockout of AKG transport between the cytosol and mitochondria has a negative impact during EMT, providing clues about cellular compartment dynamics and their impact on cancer metastasis. We hypothesize that accumulation of AKG within the mitochondria counteracts oncometabolite effects through additional regulatory mechanisms.

Two other reactions in central carbon metabolism had high priority scores, namely, enolase (ENO) and lactate dehydrogenase (LDH_L), both predicted from the same EMT proteomics data (Keshamouni et al. 2006). Upregulated enolase levels are associated with promoting cell growth, migration, and invasion during EMT in various cancers (Song et al. 2014; Zhao et al. 2015). Further, LDH is highly associated with cancer metastasis, and has been shown to activate EMT in several cancers, including lung cancer, during metastasis (Hou et al. 2019; Zhang et al. 2018). While it is known that ENO and LDH upregulation and/or increased activity are associated with poor patient prognosis, little is known about how changes in metabolic activity over time leads to differential sensitivity in cancer. Our modeling approach shows that enolase and lactate dehydrogenase are essential during later stages of EMT compared to earlier stages, revealing information about time-dependent sensitivity of these well-known targets for the first time.

In addition to nutrient exchange reactions, we found three metabolic reactions that consistently decreased growth upon KO across all time points. Two metabolic enzymes were involved in fatty acid metabolism: Fatty acid CoA ligase hexadecanoate and beta-ketoacyl synthetase. Fatty acid synthase (FASN) is a potential therapeutic target for NSCLC, and beta-ketoacyl synthetase is one component of FASN. Preclinical studies show that beta-ketoacyl synthase inhibition induces apoptosis and stops proliferation in cancer cells *in vitro* and *in vivo* (Menendez et al. 2004; Pizer et al. 1996).

Further, our model suggests two additional reactions that do not have literature backing to be potential therapeutic targets, but are associated with metabolic pathways that are frequently dysregulated across different cancers. Lipid metabolites concentrations have prognostic value, and dysregulated fatty acid metabolism is associated with poor cancer patient prognosis. While it is known that metabolic enzymes such as Fatty Acid CoA Ligases modify ratios of these fatty acids, and that there is differential regulation and expression of these metabolic enzymes in cancer, little is known about the balance of fatty acids and fatty acyl-CoAs and its impact on cancer. Our model suggests that the fatty acid CoA ligase that specifically modifies hexadecanoate contributes highly to EMT and suggests an interesting hypothesis that needs to be validated experimentally, with potential for therapeutic intervention.

Additionally, we analyzed reactions that either had very strong effects on a single study or were predicted to impact biomass in specific time points in at least two out of five studies (**S. Figure 2**). This provided us information about reactions that show temporal-specificity or robustness across datasets. Glucose and aspartate exchange reactions were predicted to be sensitive across all time points and experiments, suggesting that cells in all stages of EMT are sensitive to perturbations to these nutrients. It is well documented in the literature that high glucose levels facilitate migration and invasion processes in EMT for several types of cancer (Xu et al. 2019; Liu et al. 2016). Additionally, aspartate is crucial to cell proliferation and survival in cancer (Birsoy et al. 2015; Alkan et al. 2018; Sullivan et al. 2015). Our model also captured aspects of metabolic heterogeneity associated with glutamine metabolism in EMT. We found that cells were dependent on glutamine exchange in early (1 hr) and late (48-72 hrs) time points, while becoming insensitive to glutamine exchange during intermediate stages (8 - 24 hrs) (**S. Figure 2; top row**). Glutamine metabolism is essential for sustaining proliferation in many tumor lineages including NSCLC, and the dysregulation of glutaminolysis is a hallmark of cancer metabolism (Yang et al. 2017). Glutamine regulates the activation of STAT3, a critical transcription factor associated with tumor growth and metastasis (Cacace et al. 2017; Yang et al. 2014). Together, these results suggest that our COBRA models can accurately predict well-known impact of nutrient perturbations in cancer and EMT.

### Isoform-specific role of Enolase 3 in regulating cell survival during EMT

Reactions in glycolysis, especially enolase, was identified by both our flux and gene knockout analysis to have high metabolic activity and sensitivity to knockout. During tumor progression, cancer cells must increase glucose metabolism. Owing to the hypoxic tumor microenvironments, cancer cells upregulate glycolytic enzymes, including Enolase (Eno), to support anaerobic proliferation (Warburg effect). Enolase (Eno) is a key glycolytic enzyme that catalyzes the dehydration of 2-phosphoglycerate to phosphoenolpyruvate. It occurs as 3 isoforms, Eno1 (ubiquitously expressed in all cells), Eno2 (neuronal specific) and Eno3 (muscle specific) (Chang et al., 2006). Our transcriptomic analysis show that Eno3 the muscle specific isoform which is catalytically more efficient, is 10 fold differentially expressed in cells undergoing EMT (**S. Figure 3**) (Keshamouni et al., 2006). siRNA mediated inhibition of Eno3 selectively induced apoptosis in cells undergoing EMT whereas, inhibition of the ubiquitously expressed isoform, Eno1, did not, as assessed by Annexin-V/PI staining by flow cytometry (**Figure 3A**) and Caspase8 activation assay (**Figure 3B**). These observations suggest that EMT induces reprogramming of glycolysis to an Eno3 dependent pathway to meet the energy demands of migratory and invasive cells. Inhibition of Eno3 will selectively kill cells undergoing EMT and may prevent metastasis.

**Figure 3.**
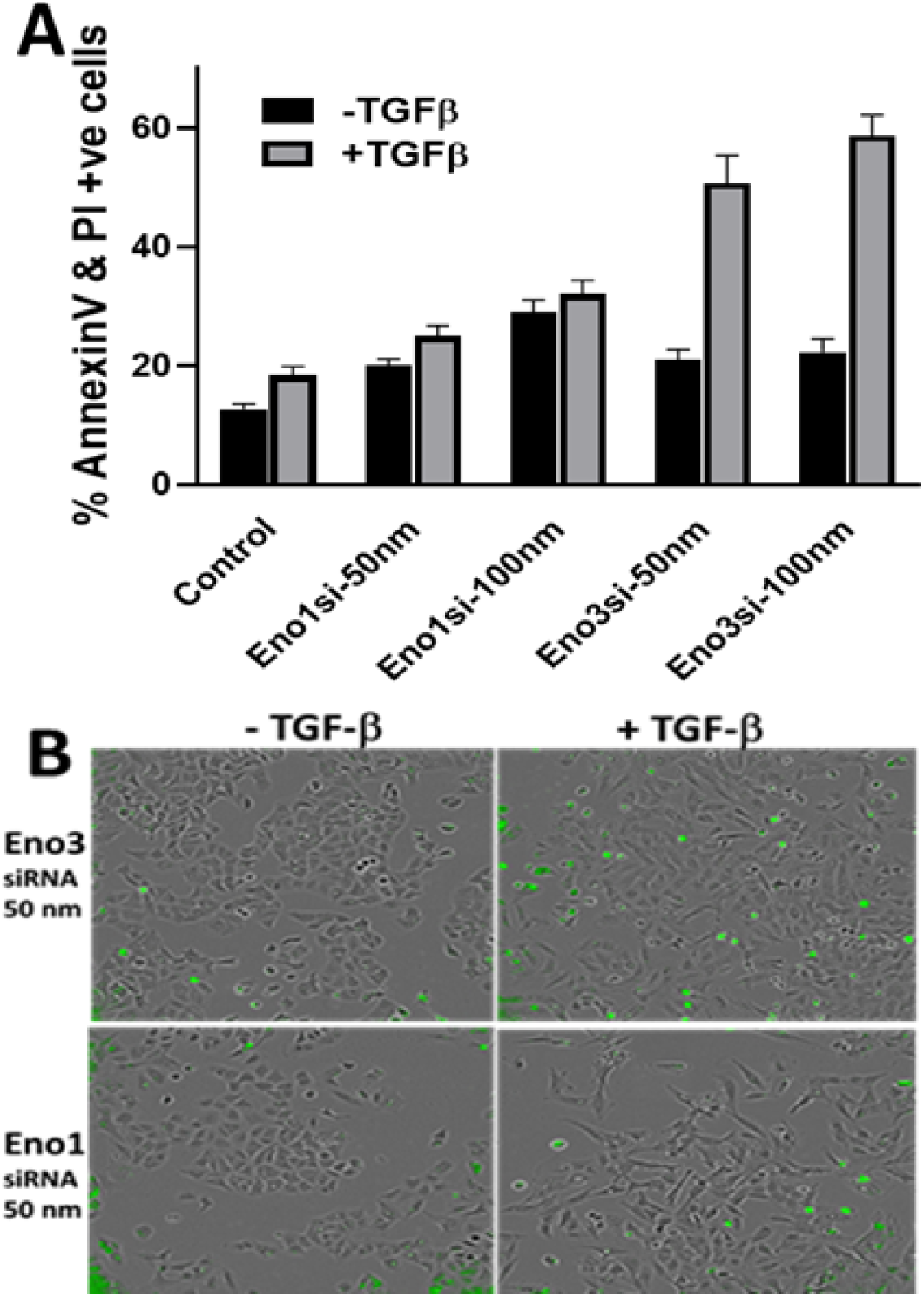
siRNA-mediated inhibition of Eno3, but not Eno1, triggers apoptosis in A549 cells undergoing EMT after 72 h TGF-b treatment. A. Apoptosis is assessed by the percentage of AnnexinV & PI positive cells by Flow cytometry. B. Caspase activation is measured using a caspase8 specific substrate that fluoresces after caspase 8-mediated cleavage.

### Single-cell knockout simulations reveal metabolic heterogeneity in a cell population undergoing EMT

To determine whether the variations observed in bulk dataset analysis are true reflection of metabolic phenotypes at the single cell level, we next analyzed single cell transcriptomics data of A549 cells induced with TGF**-β**(GSE147405; Cook & Vanderhyden, 2020). To capture subtle metabolic differences as cells’ transition from E to M states, we reconstructed separate models for each cell based on its transcriptomic profile measured in the dataset. To ensure we were observing the transition between E to M in this dataset, we visualized VIM and CDH1 expression levels in the UMAP embedding and found that the expression profiles are consistent with what is observed in the literature (**Supplementary Figure 4**). We used the resulting models for 644 individual cells across all time points and computed growth rates after genome-scale reaction knockouts in each individual cell. For comparison with the bulk datasets, we aggregated the cells, taking the average knockout scores across each time point (**Figure 4; S. Table 4)**. We found that many reactions predicted to impact growth in the single-cell analysis were also sensitive in the bulk analysis (**Table 1, S. Table 5;** N = 95 intersected reactions).

**Table 1.**
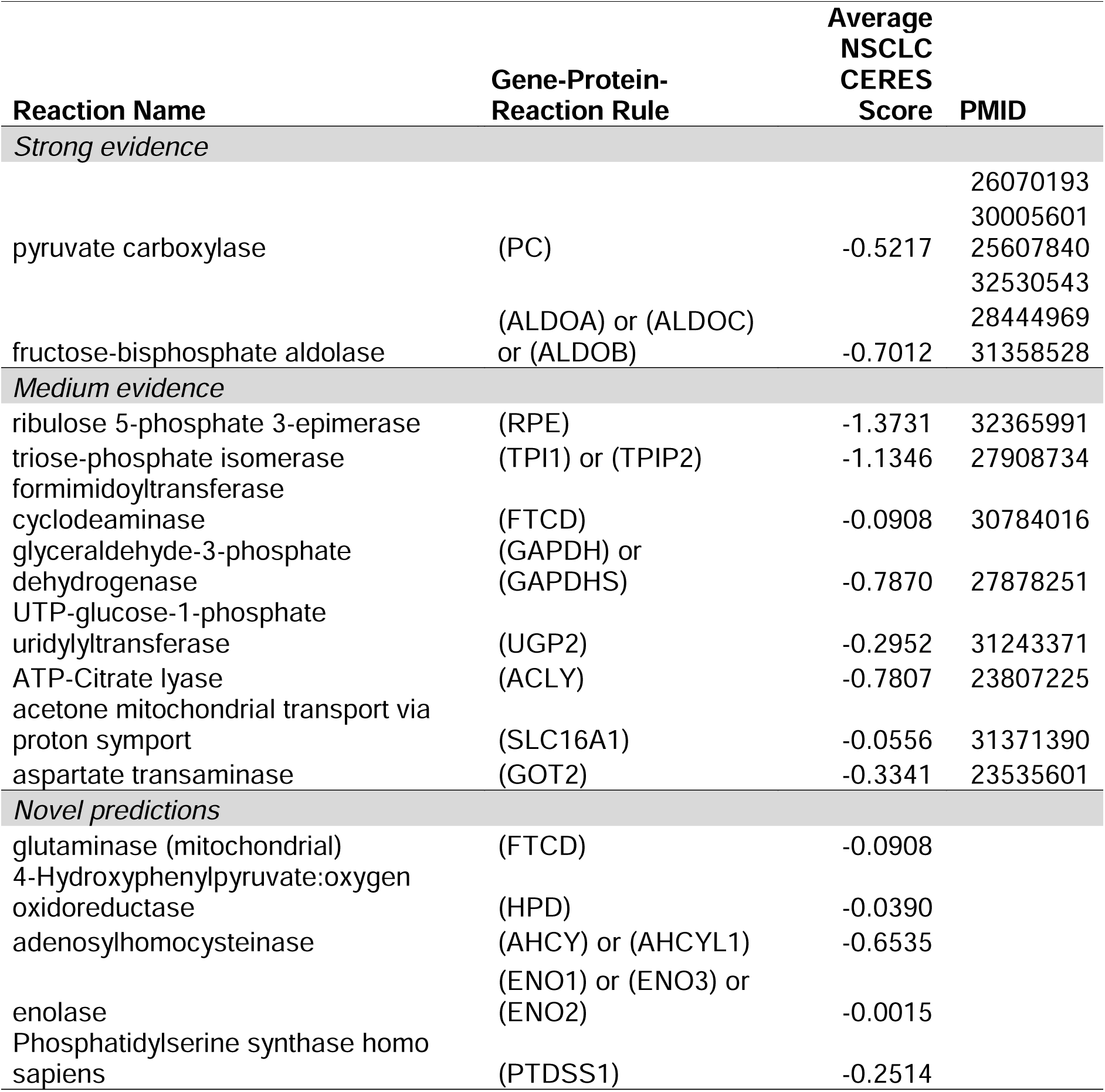
Literature review of known and novel essential reactions predicted from both bulk and single-cell simulations. For full list of all predicted essential reactions, see **S. Table 5**.

**Figure 4.**
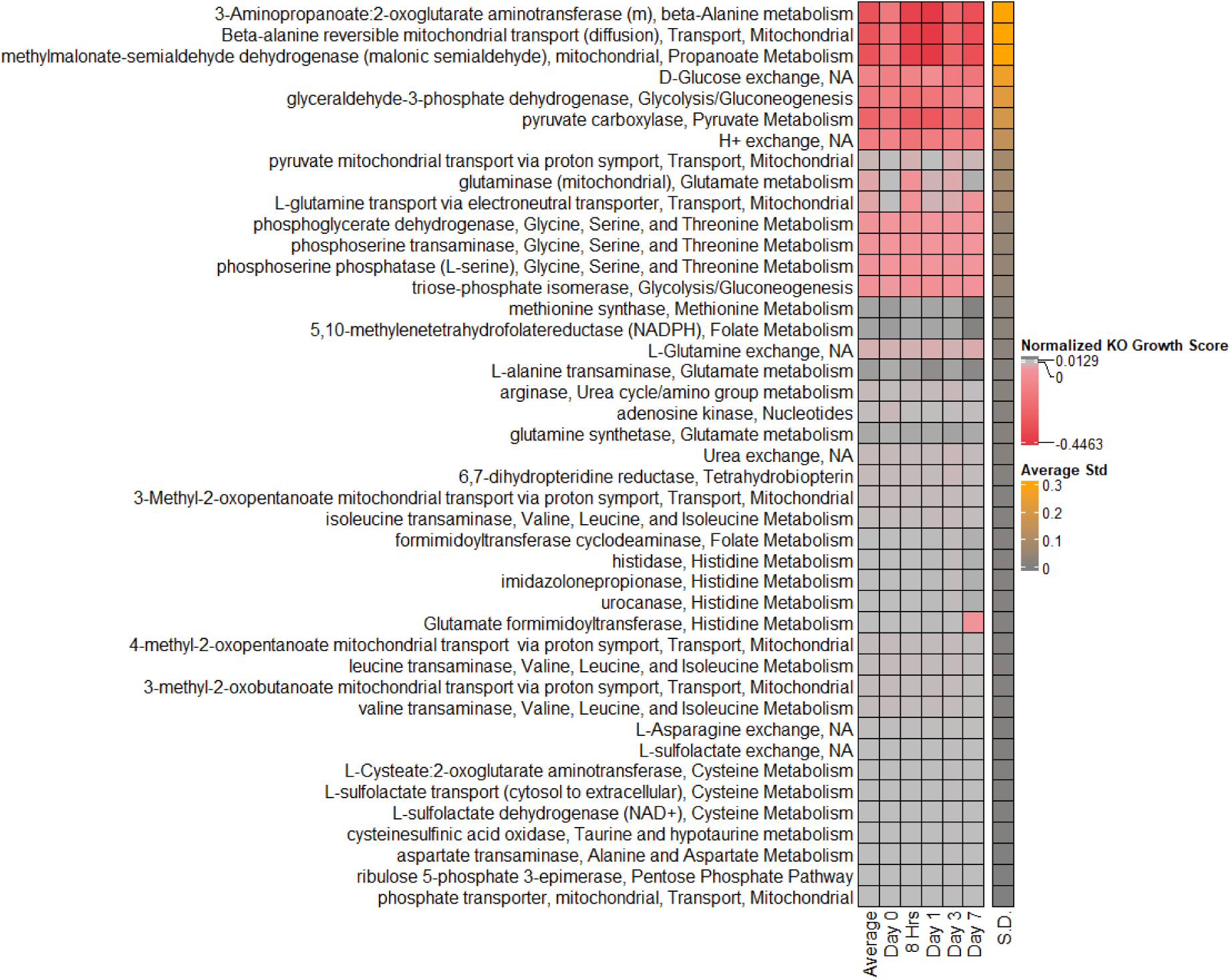
Single-cell COBRA reaction knockout analysis is consistent with results from bulk studies. **R**eaction knockout growth rates for the top 40 most variable reactions are shown in the heatmap. Column1 shows averaged data across all cells in the single-cell simulations, and data from cells grouped by time points are shown in subsequent columns (day 0, 8 hrs, day 1, day 3 and day 7). This list contains many reactions that were also found to be sensitive upon knockout in bulk studies (Table 1).

To examine how reaction knockout sensitivity changes over EMT progression for the top 5 variable central carbon metabolism reactions, we plotted the growth scores for representative reactions that had high variance onto the UMAP embedding (**Figure 5 A-F**). Reactions that were predicted from our bulk knockout profiles including AKG-malate transport, Enolase, Carnitine O-acetyltransferase, and ATP-Citrate Lyase show heterogeneous sensitivity across all time points. Interestingly, each of these vulnerabilities affects a distinct subpopulation of cells. Citrate Synthase was predicted to be sensitive across all time points, suggesting that this reaction is critical in all stages of EMT. We also observed metabolic heterogeneity in glutamine metabolism. We found that there was a positive correlation between the master regulator STAT3, glutamine synthetase, and glutamine transporter levels in the single cell data (**S. Figure 5A & B**). This is consistent with studies that have observed that glutamine regulates the activation of STAT3. Overall, our model identifies individual cells that are sensitive to specific reaction knockouts, providing a granular metabolic dependency profile of a population of cells undergoing EMT.

**Figure 5.**
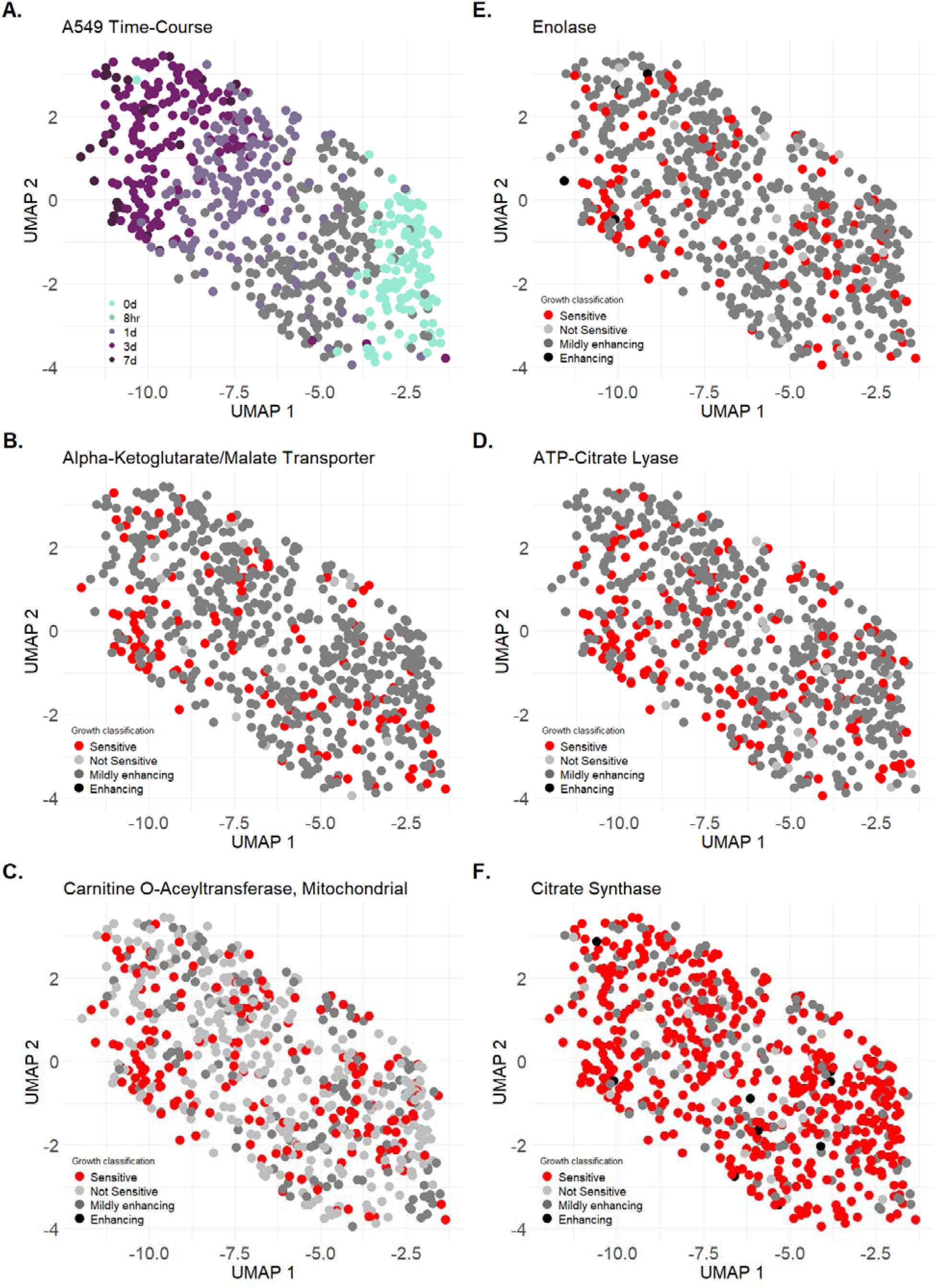
Visualization of Single-cell COBRA reaction knockout data. **A**. Shown is the UMAP visualization of the temporal trajectory of cells induced with TGF-B. **B – F**. Single-cell knockout growth rates were overlaid onto a UMAP embedding for A549 TGF-B single cell data. Growth rates were scaled from 0 to 1, where 1 indicates no change in growth rate between the cancer cell and control, while 0 indicates cell death in the cancer cell relative to control. Selected single-cell growth rate profiles for reactions that were sensitive in bulk reaction knockout simulations are shown in B-E. Growth scores (g.s.) were discretized into sensitive (g.s. < 0; red), not sensitive (g.s. = 0; light gray), mildly enhancing (0 < g.s. < 0.3; dark gray) and enhancing (g.s. > 0.3; black). The citrate synthase reaction (panel F) was selected as a control as it is an essential metabolic reaction, and it correctly shows sensitivity across all time points.

### Prioritization of metabolic targets during EMT using both bulk and single-cell simulations

We performed extensive literature curation for genes that were found to show growth reduction upon knockout in both the bulk and single-cell analysis (**Table 1, S. Table 5;** N = 95 intersected reactions). These reactions were prioritized based on the number of studies found for each gene query and its relevance to cancer and EMT. Two high confidence predictions that were found in both analyses included Pyruvate Carboxylase and Fructose-Bisphosphate Aldolase, which were shown to contribute specifically to NSCLC progression and metastasis (Sellers et al. 2015; Fu et al. 2020). Eight reaction predictions had some evidence of being dysregulated in another cancer subtype, but not NSCLC. The remaining 10 reactions have not been highlighted in the literature, and present opportunities for experimental validation.

To further assess our model predictions against experimental data, we compared our bulk and single-cell knockout results against batch-corrected CRISPR-Cas9 essentiality knockout screens integrated from the Broad and Sanger Institute (Pacini et al. 2021). Given the limited availability of CRISPR-Cas9 screenings in EMT studies, we took NSCLC cancer cell lines from the DepMap dataset, which annotated them to be derived either from a primary tumor or a metastatic site. Further, we took EMT signatures from MSigDB and EMTome to classify cancer cell lines from the cancer cell line encyclopedia (CCLE) into epithelial-like or mesenchymal-like cell-lines (**Methods**). When comparing the classification of CCLE cancer annotation with our EMT classification, we found that there was high agreement between cancer cell lines obtained from a primary site and the epithelial state while there was low agreement between the mesenchymal cell state and cell lines from a metastatic site **(Figure 6A)**.

**Figure 6.**
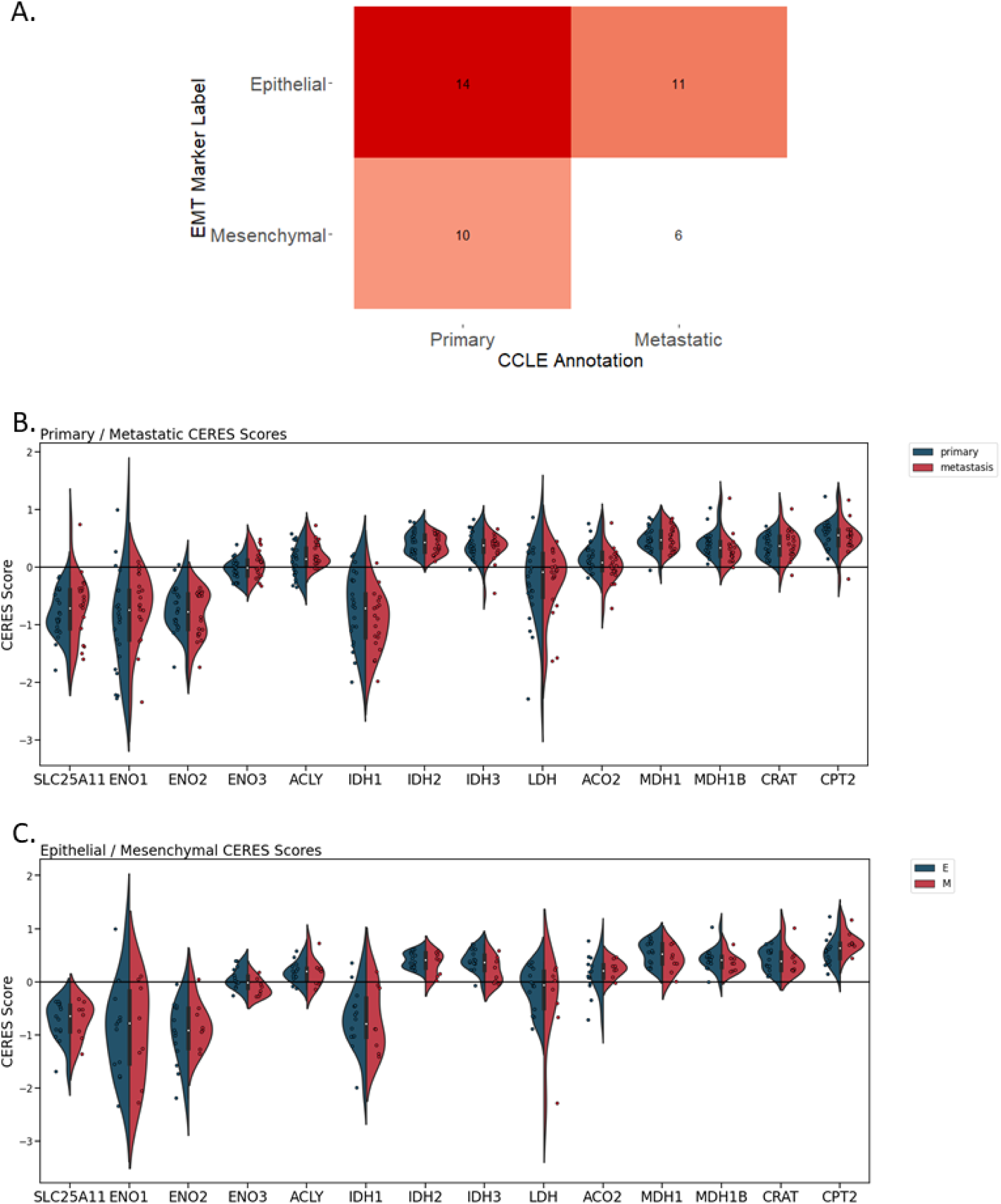
COBRA-prioritized NSCLC CERES Scores reveal metabolic vulnerabilities during EMT. A. Frequency matrix for comparing the Cancer Cell Line Encyclopedia (CCLE) primary/metastatic annotations against our Epithelial/Mesenchymal annotations based on MSigDB and EMTome signatures. Cells from the CCLE were classified as epithelial or mesenchymal based on the number of genes were up/downregulated that matched the MSigDB/EMTome signatures (**Methods**). B. Reactions predicted by COBRA to be sensitive were compared against CERES Scores. Cell lines were classified as primary or metastatic, and their distributions are shown on the violin plots. Overall, the average predicted growth scores and fluxes in hour 72 across all 5 experiments agreed with the Primary / Metastatic CERES Scores Ratios (R = 0.31 and 0.2; P-value = 0.005 and 0.035 respectively; **S. Table 6 and 7)**. These 10 metabolic genes were selected based on reactions of interest from our bulk COBRA knockout profiles and single-cell flux profiles (from **Figure 2**). C. The same analysis was repeated with the epithelial/mesenchymal annotations for sensitive reactions. The predicted growth scores and fluxes from the single-cell simulations (GSE147405) agreed with the Epithelial / Mesenchymal CERES Scores Ratios (R = 0.28, p-value=0.01).

The essentiality of metabolic enzymes identified from our model predictions were interpreted using the CRISPR gene knockout (CERES) Scores, where a lower score is associated with a higher likelihood that a given gene is essential for survival in a given cell line (Meyers et al, 2017). A score of 0 was used as the threshold to indicate the median effect of non-sensitive genes. We overlaid the CERES Scores for metabolic genes corresponding to reactions predicted from our sensitivity analysis with the CCLE cancer cell line annotation and our EMT annotation (**Figure 6B and C**). Overall, we found that the median values for both classification methods agreed with each other for the most part. The alpha-ketoglutarate / malate transporter SLC25A11, ENO1, ENO2, ENO3, IDH1, and LDH show lower median CERES scores than the threshold of 0, supporting our model’s findings. From this analysis, we were able to identify isoform-specific sensitivity in NSCLC, analogous to our validation of Eno3 dependency (**Figure 3**). IDH1 gene depletion is associated with NSCLC essentiality, compared to IDH2 and IDH3 depletion, suggesting that targeting IDH1 expression in NSCLC may be an effective therapy to supplement existing therapeutics that target specific IDH1/2 mutations.

To evaluate how well our knockout growth score predictions performed against CRISPR-Cas9 experimental data, we took the ratio of the CERES scores for the Primary site derived NSCLC cell lines to the scores for the Metastatic site derived cell lines for each metabolic gene. We found that the average growth scores and metabolic fluxes agreed significantly with the ratio data with R = 0.31 and 0.20 respectively, P-value = 0.005 and 0.036 (**S.Table 6 & 7**). We determined the correlation between our predictions against all cell lines as well, but found that the correlations were not significant with the pan-cancer CERES Score data. These results match our expectations, as our COBRA models were constrained using A549 transcriptomics and proteomics data. We evaluated the quality of each dataset on COBRA predictions, and found that the single-cell RNASeq data best matched the CERES Score ratios (KO R = 0.27, KO P-value = 0.01; Flux R = 0.20, Flux P-value = 0.03; **S. Table 6 & 7**) while bulk transcriptomics and proteomics data were weakly correlated (R < 0.1; P-value > 0.05; all bulk experiments).

There are several confounding variables that could contribute to the reduced correlation between our predictions and the experimental data. First, we assumed that cells derived from primary tumor sites have similar metabolic attributes to epithelial-like cells, while metastatic cell lines were similar to mesenchymal cells. We addressed this assumption by re-classifying cells based on EMT gene markers. We obtained similar correlations and p-values from grouping cell lines either using the CCLE annotation or our EMT classification method (**S. Table 8 & 9**).

While the transcriptomics and proteomics EMT datasets to build the metabolic models were induced using TGF**-β**, the cancer cell lines used in the CRISPR screen were not induced with an EMT inducer. Thus, the cell dynamics that occur during EMT are not captured in CRISPR screening data. Finally, we used NSCLC CCLs to evaluate the statistical significance of our results, while our simulations were performed on A549 exclusively. Due to these confounding factors, in contrast to our siRNA knockdown experiment, we found that the CERES scores did not distinguish between the three isoforms of enolase. Despite these assumptions and considerations, we found that our simulations were correlated significantly with the CERES Scores, suggesting that our model is able to extract relevant biological insights.

## Discussion

Here we utilize constraint-based modeling informed by multiple omics data sources to predict metabolic activity and knockout sensitivity during EMT. Our predictions are supported from literature validation, siRNA knockout studies, and CRISPR-Cas9 essentiality panels. We further provide a list of high confidence metabolic reaction dependencies during EMT for future experimental validation. Our approach also provides insights into metabolic activity at the single-cell level, which is not possible to infer with current experimental methodologies.

Our modeling identified metabolic enzymes that are novel as well as those with experimental evidence in literature supporting their role in tumor progression. We identified known metabolic reactions that contribute to cancer progression, such as glucose and glutamine transport. We further identified metabolic reactions associated with fatty acid metabolism that contribute to metastasis. We found that most glycolytic reactions were overactive in the early and late stages of EMT. This time-dependent aspect of glycolytic activity was intriguing and suggests a potential vulnerability during EMT. We further experimentally validate the essential role of the enolase reaction in EMT. The enolase enzyme is implicated in cancer progression for various tissue lineages, but so far has not been identified as a crucial player in NSCLC metastasis. Our COBRA modeling approach identified reaction catalyzed by Enolase as highly active during the early and late stages of EMT and predicted enolase knockout to have a negative impact on cellular growth. Enolase has three isoforms with a degree of cell type specific expression. Eno1 is ubiquitously expressed in all cells, Eno2 is neuronal specific and Eno3 is a muscle specific isoform. In our transcriptomic data sets, we observed expression of Eno1 and Eno3, but not Eno2. Even though COBRA analysis did not distinguish between isoforms, we were able to experimentally demonstrate an isoform specific function for Eno3 in cell survival during EMT. This is consistent with the kinetically more active muscle specific Eno3 regulating energy-intensive migratory behavior of cancer cells. This observation also fits with the broader trend between catalytic activity and over-expression observed across various cancers (Oruganty et al. 2020).

Comparison of our model predictions against CRISPR knockout gene essentiality scores from cancer cell lines revealed a significant correlation. Interestingly, single-cell knockout simulations were more correlated with CRISPR-Cas9 gene knockout essentiality data than models derived from bulk omics data. CRISPR-Cas9 essentiality screening is a promising high-throughput approach to determine the contribution of individual genes on cell viability. The correlation between our single-cell knockout simulations and CRISPR-Cas9 knockout essentiality data suggests that our model captures vulnerabilities during EMT. In addition to Eno3, we found that glutaminase (FTCD), 4-hydroxyphenylpyruvate oxidoreductase (HPD), adenosylhomocysteinase (AHCY), and phosphatidylserine synthase (PTDSS1) to be novel reactions that have no literature backing but have negative CERES Scores (i.e. impacts viability) in NSCLC cancer cell lines. The reactions prioritized by our model are strong candidates for drug development because they reduce cell growth in cells from later timepoints (mesenchymal/metastatic-like) relative to earlier ones (epithelial/benign-like). In addition, our model predicted ATP-Citrate lyase (ACLY) to be essential in mesenchymal-like cells. ACLY has been implicated as a crucial metabolic enzyme that facilitates cancer progression and its upregulation is associated with poor patient prognosis (Migita et al., 2008).

In summary, we present a computational model that captures metabolic activity and gene essentiality during EMT. Our modeling approach can be applied to study metabolism at a single-cell resolution and can capture the heterogeneity of other critical biological processes, including tissue differentiation and development of disease states.

## Data and software availability statement

Bulk transcriptomics data was obtained from GSE17708 and GSE17518. Bulk proteomics data was obtained from Keshamouni et al., 2006 and Lu et al., 2019. Single-cell EMT transcriptomics data was obtained from GSE147405.

All COBRA data and meta-analyses performed can be found in the supplementary table.

All scripts used to analyze these datasets can be found in this <u>GitHub repository</u>.

**Supplementary Figure 1.**
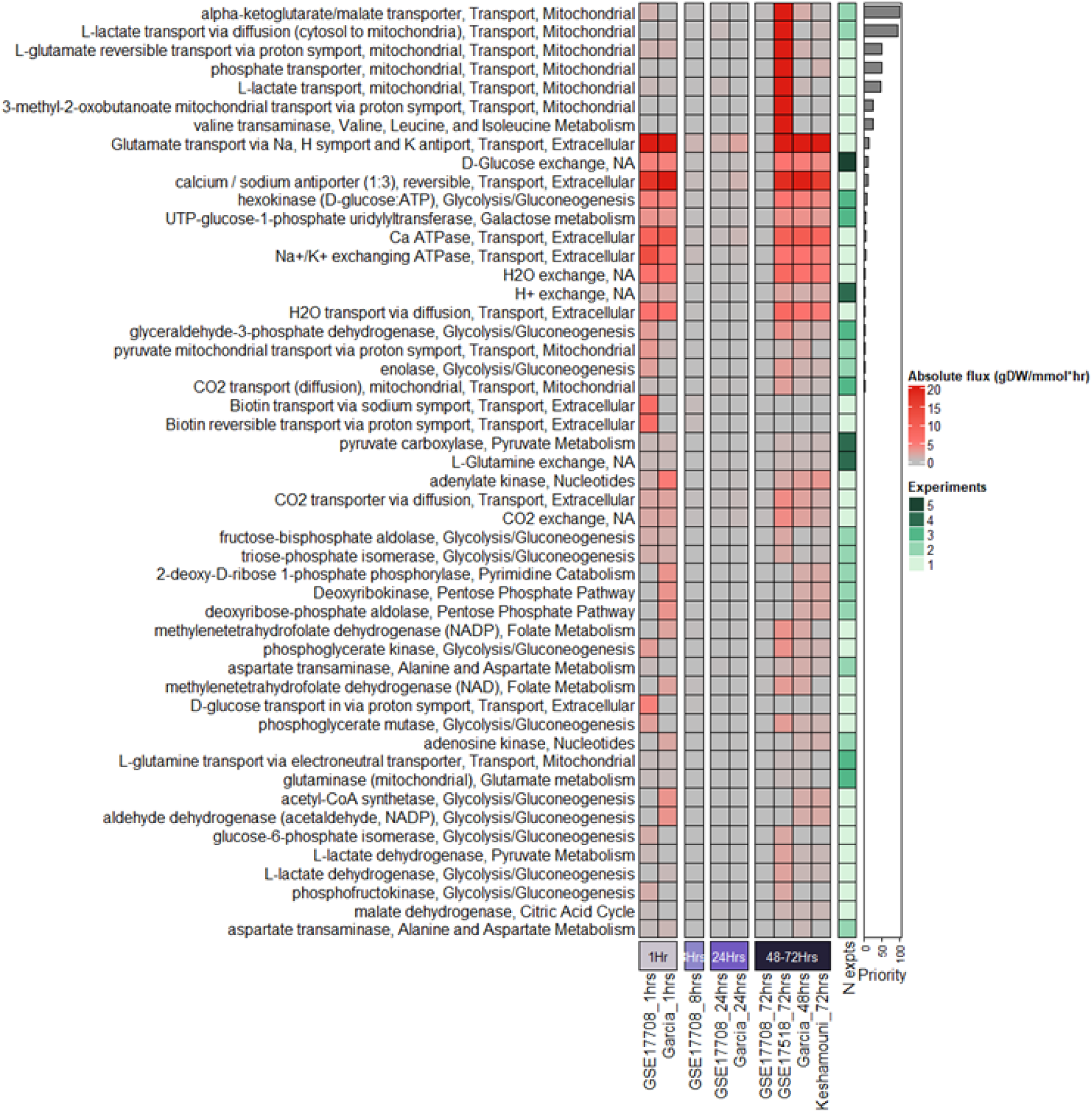
The top 50 reactions that are predicted to be overactive are ranked by priority score. (Related to figure 1). The priority score is a function of the number of studies with significant metabolic genes that encode the reaction and the effect size (Z-score or log2 fold change). The metabolic fluxes were simulated using the RECON1 metabolic reconstruction. Several EMT associated metabolic reactions predicted by our model such as Glycolysis/Gluconeogenesis, Glutamine metabolism and Nucleic acid metabolism are commonly dysregulated in cancer.

**Supplementary Figure 2.**
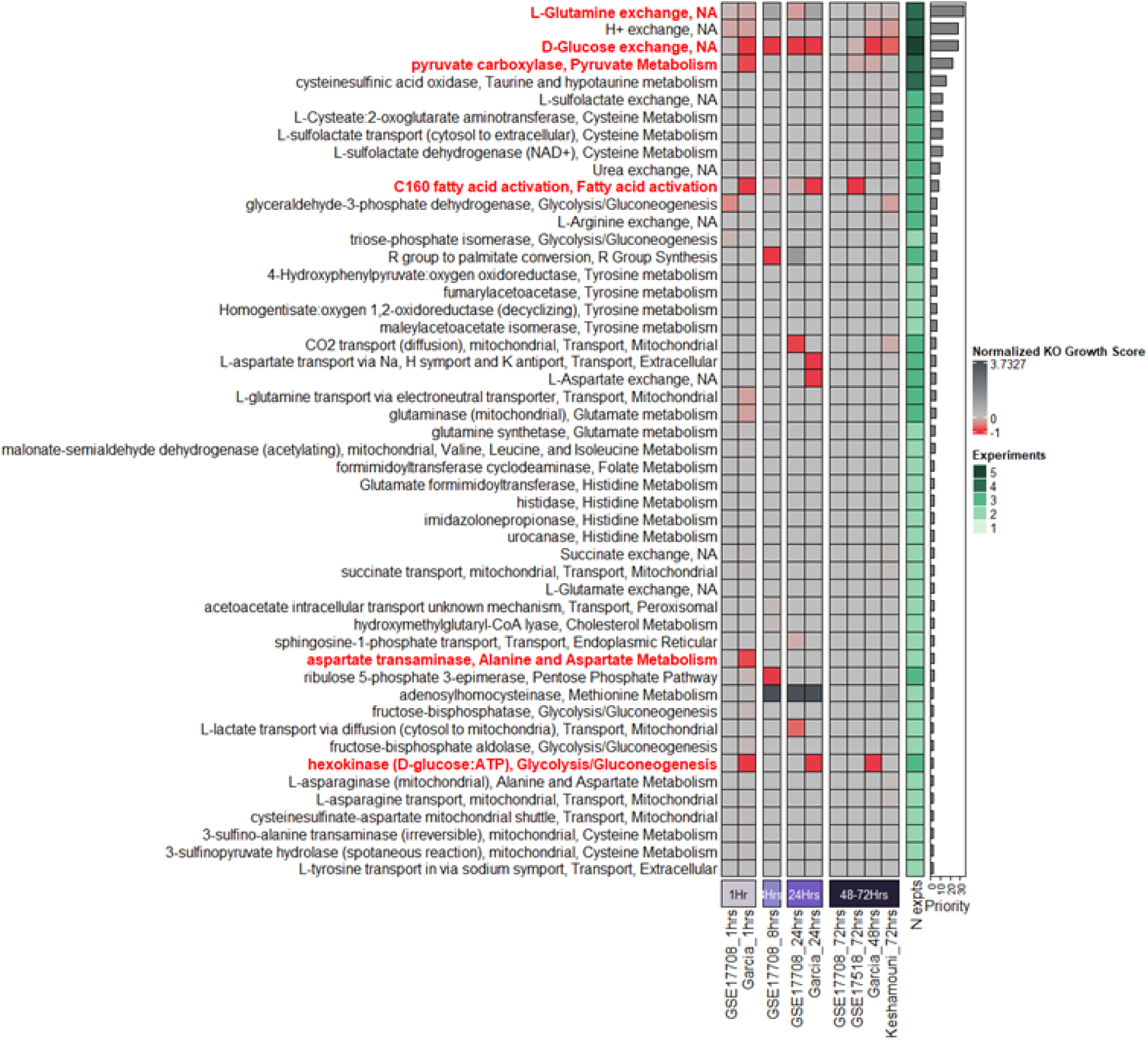
Top 50 reactions ranked by priority score predicted to be sensitive to reaction KO. (related to figure 2). The heatmap shows the top 50 reactions ranked by priority score predicted to reduce A549 growth rate upon reaction knockout in RECON1 in all bulk experiments. In contrast, Figure 2 focuses on reactions sensitive in specific EMT stages and studies, while this sensitivity profile shows the top 50 reactions by priority score. Eight nutrient exchange reactions including glucose and aspartate exchange reactions were predicted to have a negative impact on growth following knockout at different stages of EMT. Glutamine exchange interestingly decreased growth upon KO during early and late EMT timepoints but not in intermediate time-point (24hrs). Hexokinase and C100 fatty acid activation were predicted to be essential across all time points. Pyruvate carboxylase was predicted to be sensitive in early EMT (1 hour).

**Supplementary Figure 3.**
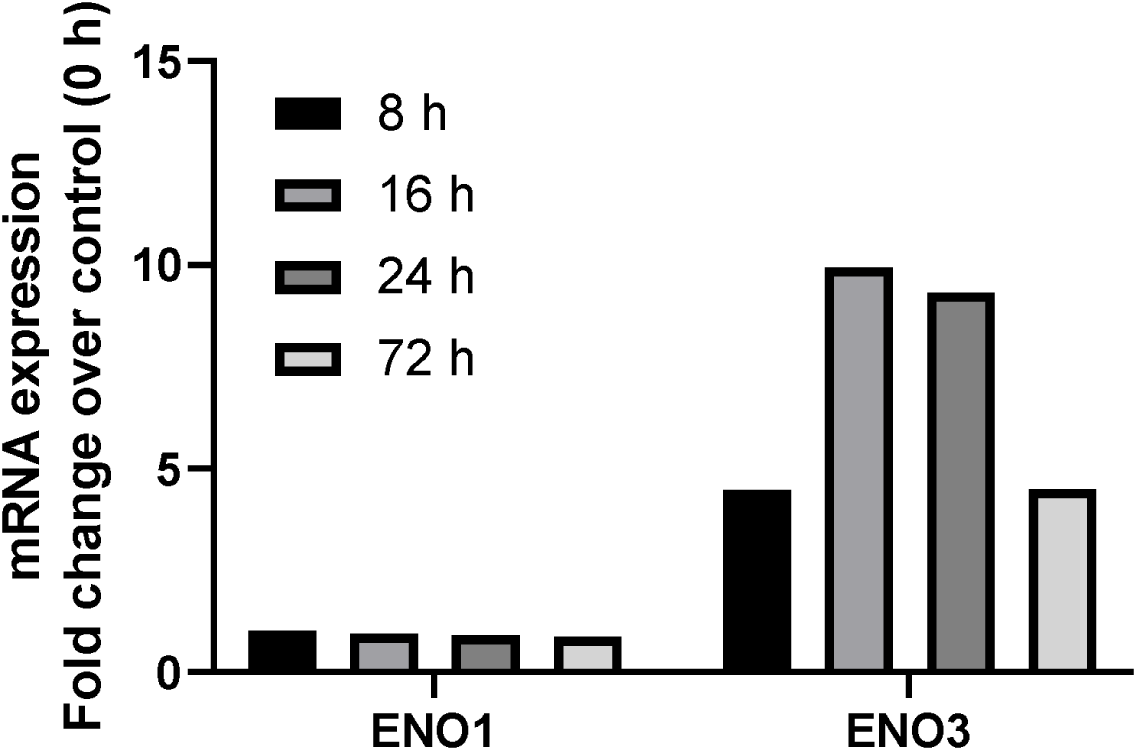
mRNA expression fold change of ENO1 versus ENO3 over time.

**Supplementary Figure 4.**
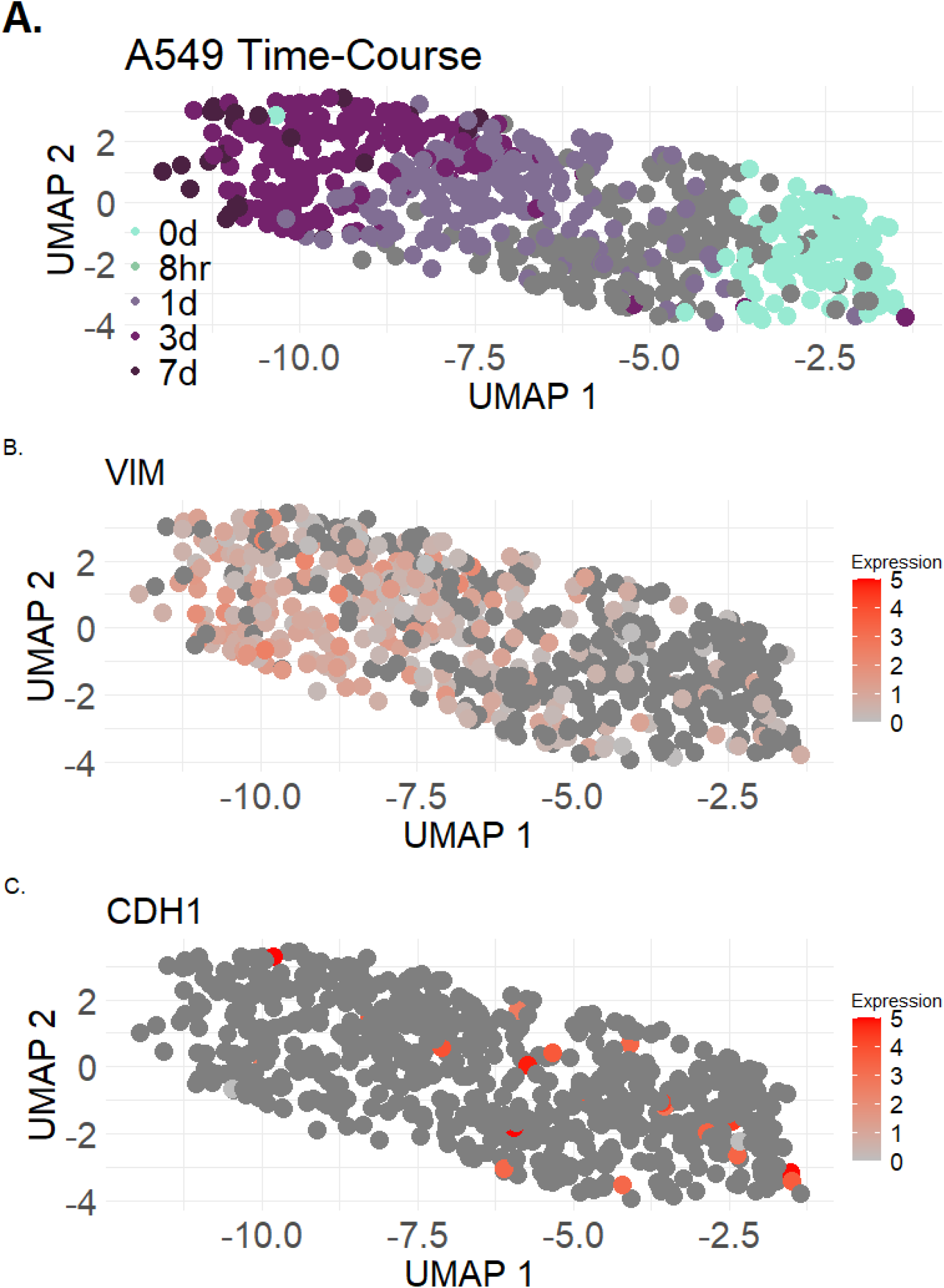
Single-cell RNASeq data of EMT biomarkers reveals cell states. A. Time-course trajectory of A549 cells induced with TGF-B. B. VIM expression over time. C. CDH1 expression over time.

**Supplementary Figure 5.**
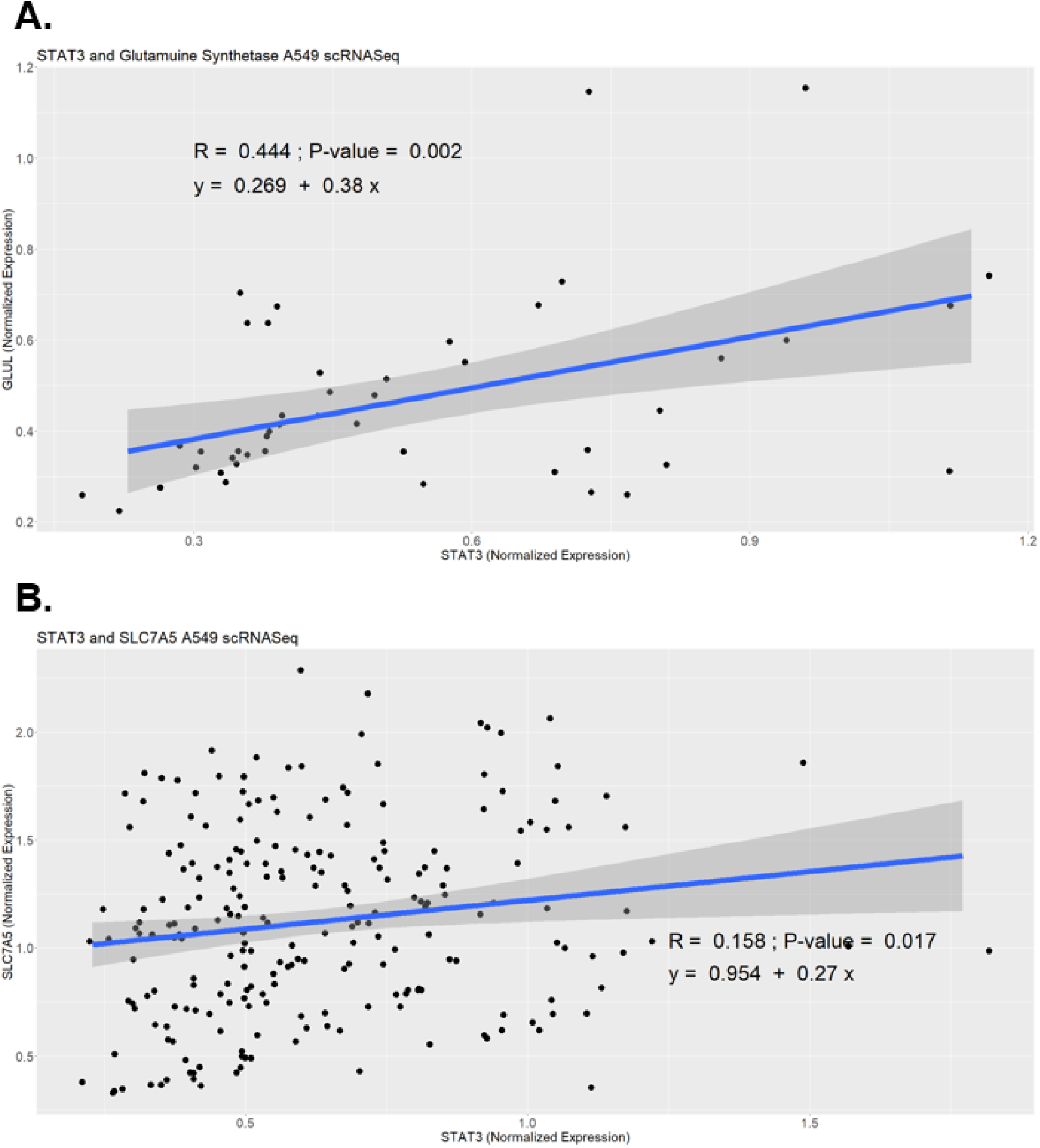
Correlation analysis between STAT3 and glutamine metabolism-related genes Glutamine Synthetase and glutamine transporter SLC7A5. A. Shown are cells that contain non-zero expression levels for both STAT3 and GLUL across cells with TGF-*β*induction. A significant positive correlation (R = 0.444; P-value = 0.002) was observed. B. Shown are cells that contain non-zero expression levels for both STAT3 and SLC7A5 across cells with TGF-*β*induction. A significant but weaker positive correlation (R = 0.158; P-value = 0.017) was observed.

